# Intronic TNR-Retained *Isopropylmalate Isomerase Large Subunit1* transcripts impair leaf development in Arabidopsis

**DOI:** 10.1101/2023.03.24.534132

**Authors:** Yimeng Li, Rui Li, Kensuke Kawade, Muneo Sato, Ayuko Kuwahara, Ryosuke Sasaki, Akira Oikawa, Hirokazu Tsukaya, Masami Yokota Hirai

## Abstract

Intronic trinucleotide repeat (TNR) is widely distributed in plant genomes. In Arabidopsis accession Bur-0, abnormally expanded TTC repeat in intron-3 of the *ISOPROPYLMALATE ISOMERASE LARGE SUBUNIT1* (*IIL1*) gene causes growth defects called the *irregularly impaired leaves* (*iil*) phenotype, triggered by DNA methylation-mediated *IIL1* gene silencing at elevated temperature. However, little is known about how the reduced expression of *IIL1* causes the *iil* phenotype. We demonstrated that the *iil* phenotype was resulted from the relative increase of intron-3-retained *IIL1* transcripts through the experiments where the *iil* phenotype was reproduced by introducing the *IIL1* gene harboring 100 copies of TTC repeat into Col-0. The *iil* phenotype appeared when the total amount of the *IIL1* transcripts was decreased by co-suppression and the percentage of intron-3-retained *IIL1* transcripts was increased. The *IIL1* gene encodes an isopropylmalate isomerase large subunit, forming heterodimers with small subunits (AtLeuD1, AtLeuD2, or AtLeuD3). In the *myb28 myb29* mutant lacking AtLeuD1 and AtLeuD2, the *iil* phenotype was almost completely suppressed regardless of higher percentage of intron-3-retained *IIL1* transcripts. The results indicated that the *iil* phenotype was associated with interaction with AtLeuDs, suggesting that intronic TNR-containing transcripts were translated into abnormal proteins and perturbed the metabolic pathway supporting the leaf development.

## Introduction

Metabolism is a fundamental process of life and closely related to development. However, how metabolism affects development is not yet fully understood. A peculiar leaf phenotype was observed by Sureshkumar *et al*. (2009), in their report on a growth defect of *Arabidopsis thaliana* (hereafter, Arabidopsis) associated with trinucleotide repeat (TNR) expansions in a metabolic enzyme. TNR expansions are a class of microsatellite that are widely distributed in the genomes of nearly all organisms (Tautz *et al*., 1986; Gur-Arie *et al*., 2000; Morgante *et al*., 2002). They are dynamic mutations, and in humans, are responsible for several neurodegenerative diseases (Pearson *et al*., 2005; Mirkin, 2007; La Spada & Taylor, 2010). In plants, TNR expansions have commonly been used as genetic markers (Varshney *et al*., 2005), but it was only recently that the effects of their expansions on plant physiology were reported. Sureshkumar *et al*. (2009) found that the Arabidopsis natural accession Bur-0 suffered from growth abnormalities and showed misshapen leaves, referred to as the “*irregularly impaired leaves*” (*iil*) phenotype, but only in elevated temperatures above 27°C. The *iil* phenotype is associated with TTC repeats that have more than 400 copies in the third intron (intron-3) of the *ISOPROPYLMALATE ISOMERASE LARGE SUBUNIT1* (*IIL1*, At4g13430, also named *AtLeuC*) allele of the Bur-0 accession (Sureshkumar *et al*., 2009) (Figure S1). The expression of *IIL1* was down-regulated by elevated temperature in Bur-0, but unaffected in Pf-0 and Col-0 accessions in which *IIL1* alleles contain only 23 copies of the TTC repeat. Consistent with the expression levels of *IIL1*, the *iil* phenotype was not observed in the Pf-0 and Col-0 accessions. Knockdowns of *IIL1* using artificial microRNA results in a more severe *iil* phenotype, while overexpression of the coding sequence (CDS) of *IIL1* restores the *iil* phenotype in Bur-0 (Sureshkumar *et al*., 2009). In the Bur-0 plants exhibiting the *iil* phenotype, a small portion of the *IIL1* transcripts retained intron-3 which contained expanded TTC repeats (Sureshkumar *et al*., 2009). Their following study demonstrated that the down-regulation of *IIL1* expression was caused by epigenetic gene silencing. In elevated temperatures, expanded TTC repeats induced the accumulation of 24-nt small interfering RNAs (siRNAs) through DICER-LIKE3, and subsequently led to DNA methylation-mediated silencing of the *IIL1* gene (Eimer *et al*., 2018). However, it is still unclear how the reduced level of the *IIL1* transcripts causes the *iil* phenotype.

The *IIL1* gene encodes the large subunit of heterodimeric isopropylmalate isomerase (IPMI). In prokaryotes and plants, IPMI is comprised of a large and small subunit (Gruer *et al*., 1997; Binder, 2010; Amorim Franco and Blanchard, 2017), and participates in the leucine biosynthetic pathway. Brassicaceae plants including Arabidopsis, which produce specialized metabolites called glucosinolates, possess a similar pathway used for the side-chain elongation of methionine. In Arabidopsis, the large subunit of IPMI is encoded by the single-copy gene *IIL1*, while the small subunit is encoded by three genes, namely *AtLeuD1*, *AtLeuD2*, and *AtLeuD3*, which contribute to substrate specificity. When IIL1 interacts with AtLeuD3, the heterodimer functions primarily as an isopropylmalate isomerase and participates in leucine biosynthesis. On the other hand, when IIL1 dimerizes with AtLeuD1 or AtleuD2, the heterodimer works as methylthioalkylmalate isomerase and is involved in methionine side-chain elongation, which precedes the biosynthesis of methionine-derived aliphatic glucosinolates (AGSLs) (Knill *et al*., 2009; Sawada *et al*., 2009; He *et al*., 2010; Imhof *et al*., 2014). Sureshkumar *et al*. (2009) demonstrated that exogenous leucine supplies did not restore the *iil* phenotype, suggesting that the changes in the leucine biosynthesis were not a primary cause of this development defect.

In this study, we aim to clarify the mechanisms underlying the occurrence of the *iil* phenotype. We generated transgenic Col-0 plants possessing the *IIL1* gene containing expanded (100 copies) TTC repeats. By using these lines, which showed the *iil* phenotype even under normal temperatures (22-23°C), we analyzed the penetrance (frequency of occurrence) of the *iil* phenotype and *IIL1* transcript levels. We revealed that the penetrance of the *iil* phenotype was dependent on the TTC repeat length. We further demonstrated that the occurrence of the *iil* phenotype was associated with an increased percentage of intron-3-retained *IIL1* transcripts in the overall total of *IIL1* transcripts. Moreover, we revealed that repression of *AtLeuD1* and *AtLeuD2* expression suppressed the *iil* phenotype. Based on these results, we propose the molecular mechanism in which intron-3-retained *IIL1* transcripts interfered with the formation of the normal IIL1-AtLeuD3 heterodimers or perturbed the function of the normal IIL1-AtLeuD3 heterodimers, resulting in the *iil* phenotype.

## Materials and Methods

### Plant materials and growth conditions

The Arabidopsis accessions Col-0 and Bur-0 were used as the WT controls. Two alleles of *IIL1* (At4g13430) knockdown mutants, *iil1-1* (*SALK_029510*, T-DNA insertion at promoter) and *iil1-2* (*SALK_065789*, T-DNA insertion to 5′-UTR), were obtained from the Arabidopsis Biological Resource Center. For phenotype observation and RT-PCR analysis, plants were grown on soil mixed with vermiculite (2:1, v/v) at 22°C under fluorescent light (a photosynthetic flux of 100 to 120 μmol photons m^-2^ s^-1^) with a light/dark cycle of 16h/8h and watered twice a week. For the analysis of the *iil* phenotypes penetrance, plants were grown on agar-solidified 1/2 Murashige and Skoog medium (Murashige & Skoog, 1962) with 1% sucrose in a growth chamber at 22°C under the same light condition. For the observation of the *iil* phenotype induced by high temperatures, plants were cultured on soil mixed with vermiculite (2:1, v/v) at either 22°C or 28°C with a light/dark cycle of 8h/16h.

### Vector construction and establishment of the transgenic lines

Genomic DNA was extracted from ten-day-old seedlings of the Bur-0 and Col-0 WT using the DNeasy plant mini kit (Qiagen, Tokyo, Japan). The *IIL1* gene containing the 5′-UTR, exons, introns, and 3′-UTR was amplified using KOD-Plus-Neo DNA polymerase (Toyobo, Osaka, Japan). The copy number of the TTC repeat within intron-3 of the amplified *IIL1* gene was 23 when the Col-0 genomic DNA was used as the template, and approximately 100 when the Bur-0 genomic DNA was used. *IIL1* amplicons were introduced into the entry vector pENTR-D-TOPO (Invitrogen, CA, USA), and subcloned into the pGWB2 vector (Nakagawa *et al*. 2007) using Gateway system (Invitrogen, CA, USA). The resulting constructs *35S_pro_:IIL1-(TTC)_100_* and *35S_pro_:IIL1-(TTC)_23_* were transformed into the GV3101 Agrobacterium strain and introduced into Col-0 WT to generate *IIL1-(TTC)_100_/Col-0* and *IIL1-(TTC)_23_/Col-0* lines, respectively (Figure 1). The *35S_pro_:IIL1-(TTC)_100_* construct was also introduced into the *myb28 myb29* double knockout mutant (*dKO*, Col-0 background) to generate *IIL1-(TTC)_100_/dKO* lines. T3 homozygous lines were selected using hygromycin resistance and subjected to further experiments. In addition, coding sequences (CDS) for the *IIL1* were amplified from Bur-0 cDNA, introduced into pENTR-D-TOPO, and subcloned into pGWB402Ω. The resulting construct *35S_pro_:IIL1-CDS* was introduced into Col-0 to create *IIL1-CDS/Col-0* lines. Two independent lines of *IIL1-(TTC)_100_/Col-0* (#2 and #4) were genetically crossed with *IIL1-CDS/Col-0* to generate *IIL1-CDS × IIL1-(TTC)_100_/Col-0* lines. Double homozygous lines were selected from the F_3_ generation by genotyping. All kits were used in accordance with the manufacturers’ instructions. The primer sequences are listed in Table S1.

**Figure 1.**
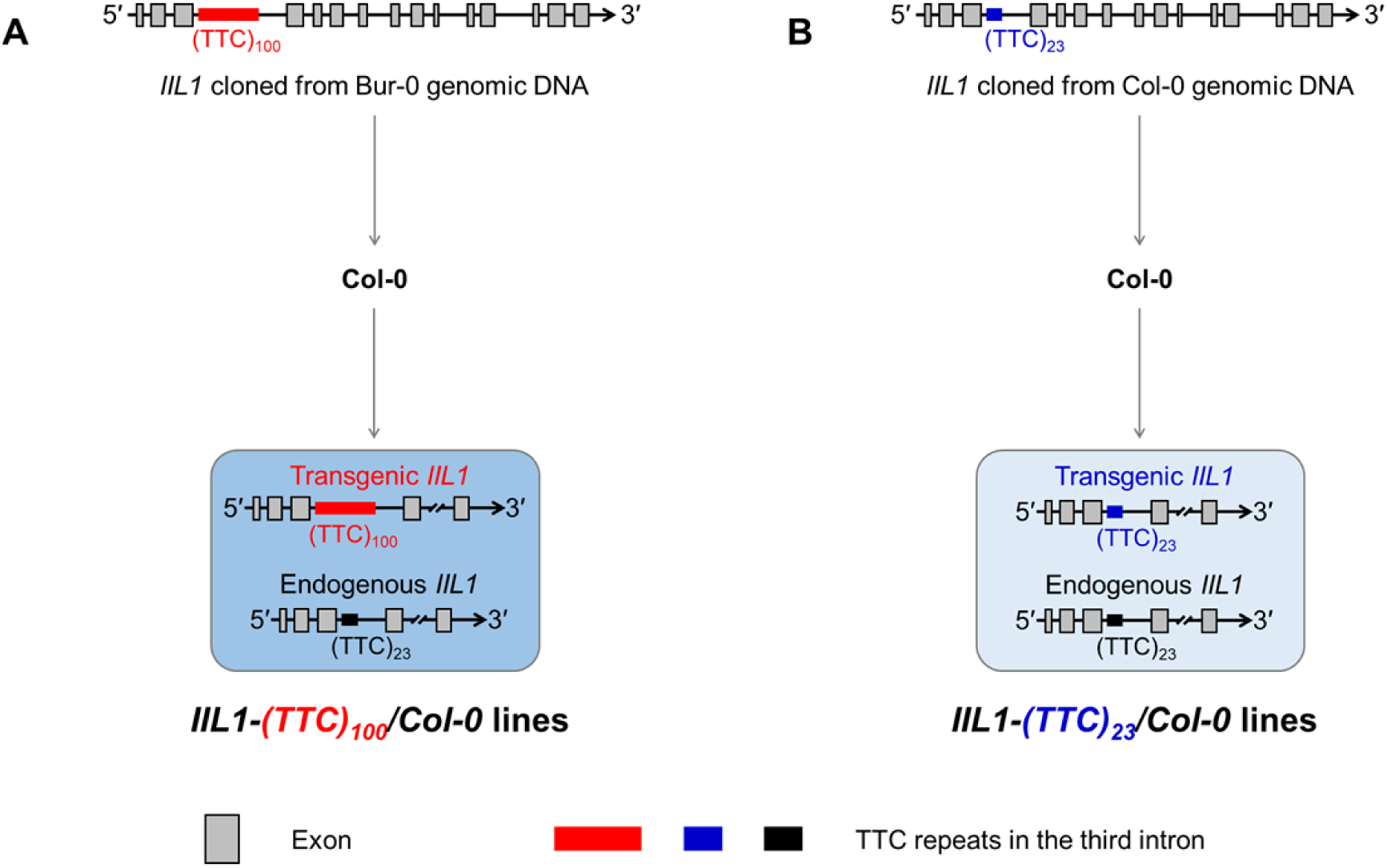
Schematic diagrams representing the construction of the *IIL1-(TTC)_100_/Col-0* and *IIL1-(TTC)_23_/Col-0* lines. The *IIL1* gene which contains approximately 100 and 23 copies of the TTC repeat in its third intron was cloned using genomic DNA extracted from the Bur-0 and Col-0 accessions, respectively. They were then introduced into Col-0 accessions to generate (A) *IIL1-(TTC)_100_/Col-0* and (B) *IIL1-(TTC)_23_/Col-0* lines, respectively. The copy numbers of the TTC repeats within the transgenic and endogenous *IIL1* genes in each line are indicated.

**Table 1.** Penetrance of the iil phenotype in IIL1-(TTC)_100_/dKO lines

| Penetrance of the *iil* |  | *IIL1-(TTC)<sub>100</sub>/dKO* |  |  |  |  |
| --- | --- | --- | --- | --- | --- | --- |
| phenotype | *dKO* | #2 | #3 | #5 | #6 | #7 |
| Stage 1.02 | 0 ± 0% | 0 ± 0% | 0.3 ± 0.5% | 0 ± 0% | 0 ± 0% | 0 ± 0% |
| Stage 1.03 | 0 ± 0% | 0 ± 0% | 0.3 ± 0.5% | 0 ± 0% | 0.3 ± 0.5% | 0 ± 0% |
| Stage 1.04 | 0 ± 0% | 0 ± 0% | 0.3 ± 0.5% | 0 ± 0% | 0.3 ± 0.5% | 0 ± 0% |
| Stage 1.05 | 0 ± 0% | 0 ± 0% | 0.3 ± 0.5% | 0 ± 0% | 0.3 ± 0.5% | 0 ± 0% |
| Stage 1.06 | 0 ± 0% | 0 ± 0% | 0.3 ± 0.5% | 0 ± 0% | 1.0 ± 1.0% | 0 ± 0% |
| Stage 1.07 | 0 ± 0% | 0 ± 0% | 0.3 ± 0.5% | 0 ± 0% | 1.3 ± 1.5% | 0 ± 0% |
| Stage 1.08 | 0 ± 0% | 0 ± 0% | 0.3 ± 0.5% | 0 ± 0% | 1.3 ± 1.5% | 0 ± 0% |

*dKO*, *myb28 myb29* double knockout mutant.

### Phenotype observations

Representative leaves were collected from four-week-old rosettes of *IIL1-(TTC)_100_/Col-0* and Col-0 WT and observed using a digital camera (Canon PowerShot S90; Canon) and a stereoscopic microscope (MZ10F; Leica Microsystems). The *iil* leaves from *IIL1-(TTC)_100_/Col-0* and normal leaves from Col-0 WT at the same leaf stage were dehydrated in an ethanol series and embedded in Technovit 7100 (Heraeus Kulzer, Wehrheim, Germany). Resin blocks were sectioned at 7 μm with a rotary microtome (Leica Biosystems), stained on glass slides with Toluidine Blue solution for 30 seconds, and observed using a Nomarski differential interference contrast microscope (BX53; Olympus).

### Analysis of the *iil* phenotype penetrance

Seeds of transgenic and WT Arabidopsis were germinated on agar-solidified 1/2 MS medium with 1% sucrose. Eight-day-old seedlings were transferred to new agar-solidified 1/2 MS medium with 1% sucrose for better growth. Three-week-old plants were used for the penetrance analysis. The growth stage of the Arabidopsis was defined based on a previous study (Boyes *et al*., 2001). The penetrance of the *iil* phenotype was determined by recording from which leaf onward the *iil* phenotype appeared in individuals and calculating the percentage of individuals exhibiting the *iil* phenotype at each stage of leaf ontogenesis. The penetrance was calculated based on three independent experiments. For each experiment, at least 100 individuals of each line were assessed.

### RNA extraction and transcript analysis

The first, fifth, and ninth normal leaves were sampled from four-week-old Col-0 WT and individuals of *IIL1-(TTC)_100_/Col-0* which did not exhibit the *iil* phenotype. The first normal leaves, the fifth transitional *iil* leaves, and the ninth *iil* leaves were sampled from individuals of the *IIL1-(TTC)_100_/Col-0* lines that exhibited the *iil* phenotype. In addition, mature normal leaves and mature *iil* leaves were sampled from four-week-old Bur-0 cultured at 22°C and 28°C, respectively. Total RNA was extracted from these single-leaf samples using the RNeasy Plant Mini kit (Qiagen, Tokyo, Japan) and treated with DNase using the TURBO DNA-free kit (Thermo Fisher Scientific, Vilnius, Lithuania). The first-strand cDNA was synthesized using ReverTra Ace qPCR RT Master Mix with a gDNA Remover kit (Toyobo, Osaka, Japan).

To detect intron-3 retention of the *IIL1* gene in *IIL1-(TTC)_100_/Col-0* lines, the cDNA was amplified by PCR using the Ex Taq DNA Polymerase (TaKaRa, Tokyo, Japan) with primers located within exon-3 and exon-4. To verify whether genomic DNA was removed from the cDNA samples, the *ACT2* gene (At3g18780) was amplified using primers spanning intron-1 to intron-3 (Appelhagen *et al*., 2011). Reverse transcription PCR (RT-PCR) products were separated in 1.6% (w/v) agarose gels containing ethidium bromide. The splicing variant of around 600 bp in size was purified using the QIAquick gel extraction kit (Qiagen, Tokyo, Japan), subcloned into the pCR4-topo vector (Invitrogen, CA, USA), and subjected to sequencing using M13 forward and reverse primers. All kits were used in accordance with the manufacturers’ instructions.

Quantitative real-time reverse transcription-PCR (quantitative RT-PCR) was performed on the StepOnePlus real-time PCR system (Applied Biosystems) using the Fast SYBR Green Master Mix (Thermo Fisher Scientific, Vilnius, Lithuania). Forward primers in the conjunction of the 5′-UTR and exon-1, and reverse primers within exon-3 were used to detect transcript levels of *IIL1* (Figure S10A). According to a single-nucleotide polymorphism in exon-1 of *IIL1^Bur-0^* and *IIL1^Col-0^*, two allele-specific forward primers were used to distinguish transgenic (originated from the *IIL1^Bur-0^* allele) and endogenous (originated from the *IIL1^Col-0^* allele) *IIL1* transcripts in *IIL1-(TTC)_100_/Col-0* lines. The first nucleotide from the 3′ end of the forward primers was designed to be matched to either *IIL1^Bur-0^* or *IIL1^Col-0^* templates, while the second nucleotide was designed to be mismatched to both templates (Figure S10B). In addition, primers located within intron-3 and exon-4 were used to detect intron-3-retained *IIL1* transcript levels (Figure 10C). For the standard curve of quantitative RT-PCR, the *IIL1* gene containing 5′-UTR, exons, and 3′-UTR was cloned from Bur-0 and Col-0 cDNA, introduced into a pENTR-D-TOPO vector, and used as the standard for the detection of *IIL1* transcripts that originated from the *IIL1^Bur-0^* and *IIL1^Col-0^*, respectively; the *IIL1* gene containing the 5′-UTR, exons, introns, and 3′-UTR was cloned from Col-0 WT genomic DNA and introduced into a pENTR-D-TOPO vector, and used as the standard for the detection of intron-3-retained *IIL1* transcripts. Standard vectors were linearized by digestion with single restriction enzymes and prepared from 10^9^ copies/μl to 10^4^ copies/μl using 1:10 dilutions with Milli-Q water. Concentrations of *IIL1* transcripts originated from *IIL1^Bur-0^*and *IIL1^Col-0^*, and intron-3-retained *IIL1* transcripts were calculated from respective standard curves. Percentages of the intron-3-retained *IIL1* were calculated as “intron-3-retained *IIL1* transcripts / transcripts from *IIL1^Bur-0^* and *IIL1^Col-0^*” in *IIL1-(TTC)_100_/Col-0* lines, “intron-3-retained *IIL1* transcripts / transcripts from *IIL1^Col-0^*” in Col-0 WT, and “intron-3-retained *IIL1* transcripts / transcripts from *IIL1^Bur-0^*” in Bur-0 WT. Average and standard deviation of the percentage were calculated from three replicates for each type of leaf sample. Primer sequences are listed in Table S1.

## Results

### T-DNA insertional *IIL1* mutants did not have the *iil* phenotype

We investigated the morphological phenotype of two T-DNA insertional *IIL1* knockdown mutants (*iil1-1*, SALK_029510; *iil1-2*, SALK_065789) in the Col-0 genetic background, in which *IIL1* expression was reduced (Knill *et al*., 2009; Sawada *et al*., 2009). In contrast to the severe *iil* phenotype observed with the artificial microRNA-mediated *IIL1* knockdown in Bur-0 (Sureshkumar *et al*., 2009), *iil1-1* and *iil1-2* did not have the *iil* phenotype (Figure S2). The results suggested that the occurrence of the *iil* phenotype could not be directly attributed to the down-regulation of *IIL1*.

### Genetic construction of IIL1-(TTC)_100_/Col-0 and IIL1-(TTC)_23_/Col-0

Among the 96 accessions of Arabidopsis, Col-0 possesses relatively long TTC repeats, as it has 23 copies (Sureshkumar *et al*., 2009) in intron-3 of the *IIL* gene; however, Col-0 did not show the *iil* phenotype at elevated temperatures, unlike Bur-0. We considered that the 23 copies of the TTC repeat were not long enough to produce the *iil* phenotype. To clarify whether longer TTC repeats could reproduce the *iil* phenotype in the Col-0 accession background, the *IIL1* gene containing expanded TTC repeats was introduced into Col-0 plants. Specifically, the full-length *IIL1* containing the 5′-UTR, exons, introns, and 3′-UTR was amplified from the genomic DNA of Bur-0. When the *IIL1* gene was cloned from Bur-0, the copy number for the TTC repeats in intron-3 was reduced to approximately 100, most likely due to the instability of the TNR during DNA duplication (Pollard *et al*., 2007). This amplicon was subcloned into an expression vector to create a *35S_pro_:IIL1-(TTC)_100_*construct under the control of the *CaMV 35S* promoter. This construct was transformed into the Col-0 wild type (WT), and the resultant transformants were named *IIL1-(TTC)_100_/Col-0* (Figure 1A). As a control experiment, the *IIL1* gene containing 23 copies of the TTC repeat in intron-3 was cloned from Col-0 and introduced into Col-0 WT under the control of the *CaMV 35S* promoter. The resultant transformants were named *IIL1-(TTC)_23_/Col-0* (Figure 1B). There is a single-nucleotide polymorphism in exon-1 of *IIL1^Bur-0^* and *IIL1^Col-0^*, which leads to a single amino acid difference in the chloroplast transit peptide of the IIL1 protein. High expression of *IIL1* in the seedlings of these transformants was confirmed (Figure S3).

### Progressive appearance of the *iil* phenotype at normal temperature in the *IIL1-(TTC)_100_/Col-0*

Occurrence of the *iil* phenotype was analyzed in four independent *IIL1-(TTC)_100_/Col-0* lines. Interestingly, the *iil* phenotype was observed at the normal temperature of 22°C in these lines (Figure S4 and S5). This phenomenon was not specific to the Col-0 genetic background, because the *iil* phenotype was also observed at 22°C in the Bur-0 lines transformed with the *35S_pro_:IIL1-(TTC)_100_* construct (Figure S6). The use of these transgenic lines enabled us to further dissect the *iil* phenotype from possible high temperature-induced morphological and metabolic changes, such as elongated petioles and membrane damage. Overall, the rosette leaves of the *IIL1-(TTC)_100_/Col-0* lines exhibited irregular shapes with incurvatures (curved upward), and mild interveinal chlorosis (Figure 2A; Figure S7 and S8). The organization of the mesophyll cells was found to be disrupted in the cross section of the irregular leaves (Figure 2B). These observations were like those for Bur-0 cultivated at high temperatures (Sureshkumar *et al*., 2009), except that *IIL1-(TTC)_100_/Col-0* did not exhibit elongated petioles. The petiole phenotype observed in Bur-0 at 27°C was shown to be regulated by the *CRY2* gene (Sanchez-Bermejo *et al*., 2015), but not the specific characteristic of the *iil* phenotype (Figure S5). Consequently, we considered that the observed phenotype of the *IIL1-(TTC)_100_/Col-0* lines was the *iil* phenotype and defined such leaves as “*iil* leaves”. In addition, we observed that leaves with a narrow shape and moderate chlorosis at the leaf bases always appeared prior to the appearance of the *iil* leaves, and thus referred to these leaves as “transitional *iil* leaves” (Figure 2A; Figure S7B and S8).

**Figure 2.**
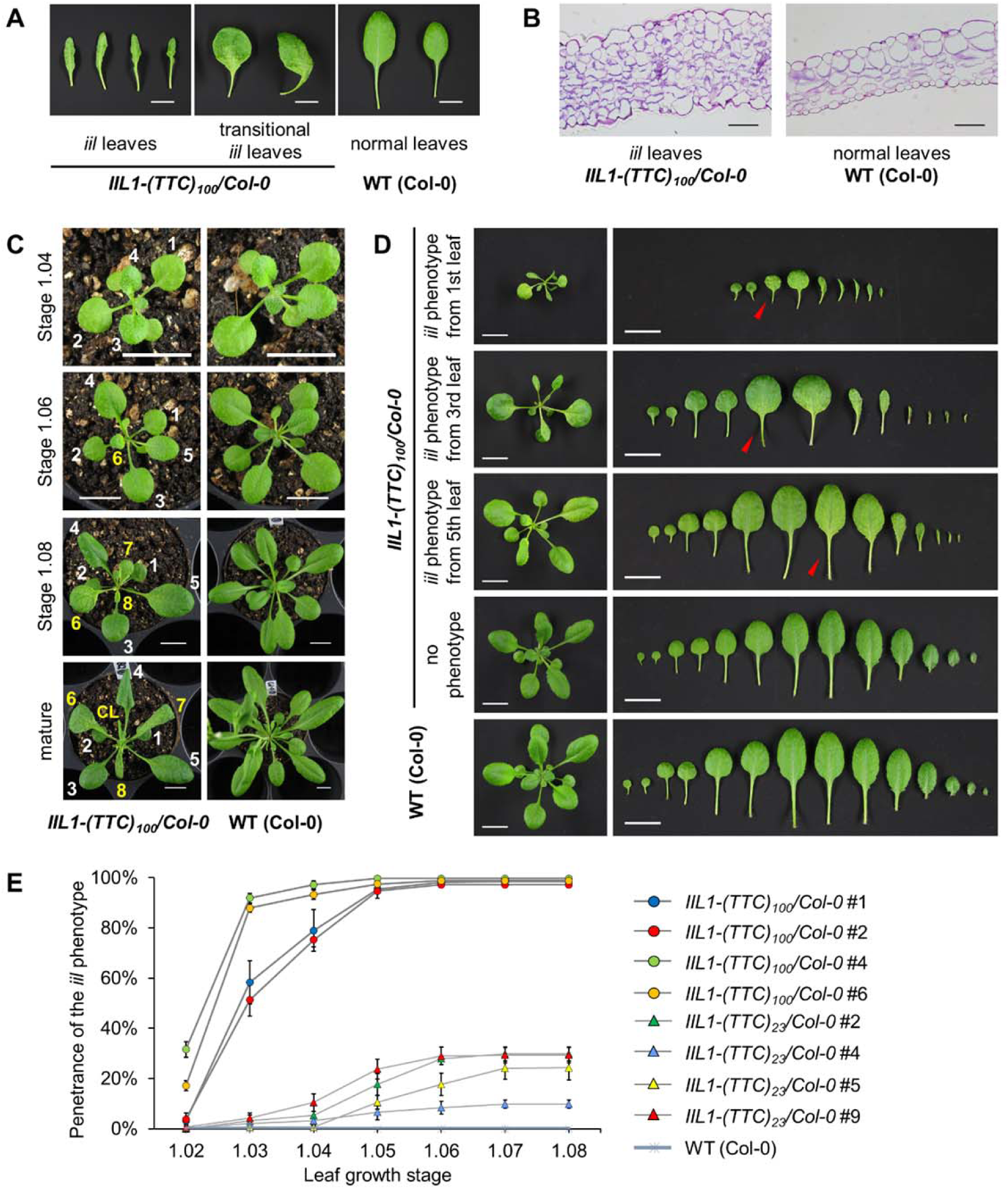
Progressive occurrence of the *iil* phenotype and its penetrance in *IIL1-(TTC)_100_/Col-0* lines. (A) Representative *iil* and transitional *iil* leaves from four-week-old *IIL1-(TTC)_100_/Col-0* plants, and normal leaves from the Col-0 wild type (WT). “Transitional *iil* leaves” were defined as abnormal leaves with a narrow shape and moderate chlorosis at the leaf base that appeared before the typical *iil* leaves. Scale bars represent 1 cm. (B) Cross section of *iil* and normal leaves from *IIL1-(TTC)_100_/Col-0* lines and the WT, respectively. The *iil* leaf showed disrupted organization of the mesophyll cells. Scale bars represent 50 μm. (C) Representative development of *IIL1-(TTC)_100_/Col-0* rosettes at the stage 1.04 (4 rosette leaves > 1 mm in length), 1.06 (6 rosette leaves > 1 mm in length), 1.08 (8 rosette leaves > 1 mm in length), and five-week-old mature plants. Corresponding WT rosettes that were the same age are also shown. The leaf number for the *IIL1-(TTC)_100_/Col-0* lines is indicated, and leaves showing the *iil* phenotype are identified with yellow numbers. CL, cauline leaf. Scale bars represent 1 cm. (D) Progressive occurrence of the *iil* phenotype at different growth stages. Representative *IIL1-(TTC)_100_/Col-0* rosettes in which the *iil* phenotype occurred from the first, third, and fifth leaves, as well as the rosette which did not show the *iil* phenotype, are presented. Red arrows indicate the leaves that developed the *iil* phenotype. Scale bars represent 1 cm. (E) The penetrance of the *iil* phenotype in *IIL1-(TTC)_100_/Col-0*, *IIL1-(TTC)_23_/Col-0* lines and the WT during leaf development. Data represent means ± standard deviations of three independent experiments. Each independent line consisted of at least 100 individuals in each experiment.

The representative development of *IIL1-(TTC)_100_/Col-0* rosettes is depicted in Figure 2C. At stage 1.04 (the development stage at which 4 rosette leaves > 1 mm in length, similarly hereafter, defined by Boyes *et al*., 2001), the shape of leaves 1 to 4 in the *IIL1-(TTC)_100_/Col-0* lines showed no obvious differences when compared to the WT. At stage 1.06, all leaves were normal in shape, but mild chlorosis was observed at the base of leaf 6 (Figure S8). At stage 1.08, a typical *iil* leaf with irregular shape, incurvature, and chlorosis (Figure 2A; Figure S7) appeared at leaf 7. In addition, leaf 6 developed into transitional *iil* leaves. When *IIL1-(TTC)_100_/Col-0* lines matured for flowering, leaves 1 to 5 remained normal, whereas the *iil* phenotype was observed in leaf 6 and in successive leaves, including the cauline leaves. Therefore, *IIL1-(TTC)_100_/Col-0* lines exhibited heterogeneous rosettes in which the earlier normal leaves and the later *iil* leaves coexisted. Such heterogeneous morphology was also observed in Bur-0 at high temperatures (Sureshkumar *et al*., 2009) or under UV-B irradiation (Tabib *et al*., 2016). In addition, the first *iil* leaves appeared at various growth stages among individuals in the *IIL1-(TTC)_100_/Col-0* lines (Figure 2d). In some individuals of the *IIL1-(TTC)_100_/Col-0* lines, the *iil* leaves did not appear in the rosettes (Figure 2d).

To quantitatively characterize the *iil* phenotype, its penetrance was investigated, which is the frequency of individuals exhibiting the *iil* phenotype at each growth stage (Boyes *et al*., 2001). As shown in Figure 2E, the *iil* phenotype appeared progressively during development. When the *IIL1-(TTC)_100_/Col-0* lines reached growth stage 1.03, more than 50% of the individuals exhibited the *iil* phenotype; while when the lines reached stage 1.08, the frequency was more than 95%. By contrast, *IIL1-(TTC)_23_/Col-0* lines, in which there were 23 copies of the TTC repeat located in the transgenic *IIL1* gene, exhibited less than 5% of *iil* penetrance at stage 1.03, and less than 35% at stage 1.08. These data revealed that the TTC repeats within intron-3 of the *IIL1* gene were responsible for the occurrence of the *iil* phenotype, and the penetrance of the *iil* phenotype was associated with the length of the TTC repeat.

### Intron-3 retention in *IIL1* transcripts in *IIL1-(TTC)_100_/Col-0*

The above-described results strongly indicate that the *ill* phenotype was not caused by loss-of-function of the *IIL1* gene. Previous studies have found that intron-3 of *IIL1* was retained in a small portion of the *IIL* transcript population of Bur-0 (Sureshkumar *et al*., 2009). To determine whether the same phenomenon occurs in *IIL1-(TTC)_100_/Col-0* lines, we analyzed the *IIL1* transcripts using reverse transcription PCR (RT-PCR).

Total RNA was extracted from the *iil* leaves of *IIL1-(TTC)_100_/Col-0* lines and normal leaves of the Col-0 WT control. The first-strand cDNA was synthesized, and the absence of the genomic DNA in the cDNA samples was verified by amplification of the *ACT2* gene using primers spanning three introns (Figure S9). To determine whether intron-3 bearing the TTC repeat was present in the *IIL1* transcripts, the cDNA was amplified using a pair of primers located within exon-3 and exon-4 of the *IIL1* gene (Figure 3A). We observed a major band slightly longer than 200 bp, which was consistent with the expected size (218 bp) of the target fragment without introns (Figure 3B). Aside from this transcript with no introns, we detected a splicing variant of around 600 bp in size specifically in the *iil* leaf cDNA (yellow arrowhead, Figure 3B). Sequencing analysis confirmed that this amplicon contained the exact boundaries of exon-3 and intron-3, and intron-3 and exon-4, as well as the intron-3 region (Figure 3C). The length of the TTC repeat in this amplicon was slightly shorter than that in the transgenic *IIL1-(TTC)_100_*gene, probably due to the instability of the TTC repeat during DNA duplication, as observed when cloning the *IIL1* gene from Bur-0. Together, these results indicated that intron-3 bearing the TTC repeat was retained in a portion of the *IIL1* transcript population in the *IIL1-(TTC)_100_/Col-0* lines (Figure 3D).

**Figure 3.**
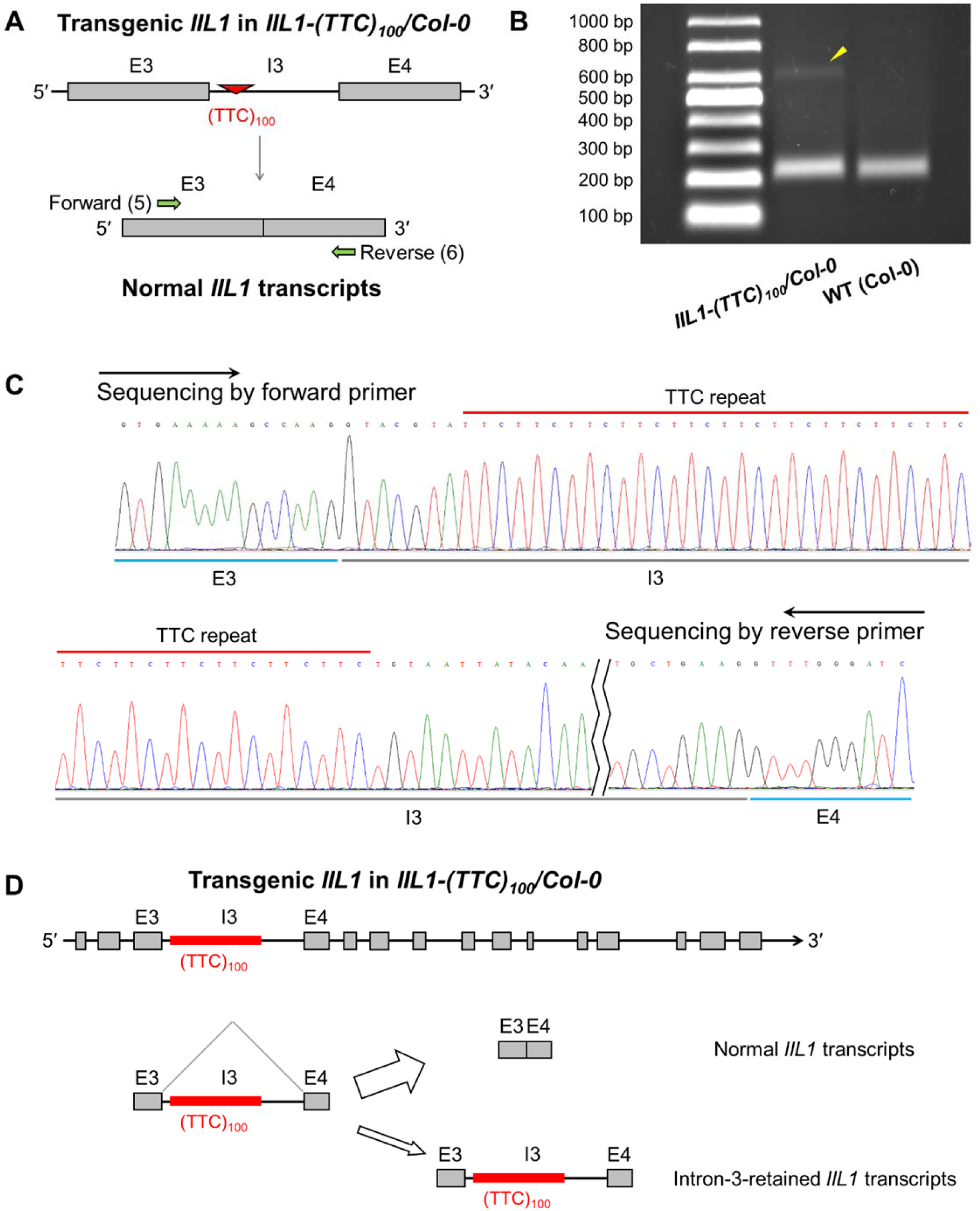
Intron-3 retention in *IIL1* transcripts of *IIL1-(TTC)_100_/Col-0* lines. (A) Localization of primers for RT-PCR analysis to detect intron-3 retention. Forward primer (serial number 5 in Table S1) and reverse primer (serial number 6) were located at exon-3 and exon-4 of the *IIL1* gene, respectively. (B) Transcription analysis of *IIL1* from the cDNA of *iil* leaves from *IIL1-(TTC)_100_/Col-0* and normal wild type (Col-0) leaves by RT-PCR using primers 5 and 6. The yellow arrowhead indicates the intron-3-retained amplicon. (C) Sequencing analysis of the intron-3-retained amplicon using forward and reverse primers. (D) Schematic diagram of intron-3 retention in the *IIL1* transcripts from the *iil* leaves of *IIL1-(TTC)_100_/Col-0* lines. When intron-3 containing the TTC repeat was spliced or retained, the *IIL1* gene was transcribed into normal or intron-3-retained *IIL1* transcripts, respectively. E3, E4, and I3 indicate exon-3, exon-4, and intron-3, respectively.

### Increased percentage of intron-3-retained *IIL1* transcripts in the *ill* leaves

We considered that intron-3-retained *IIL1* transcripts might perturb a certain process involving the normal *IIL1* transcripts and lead to the *iil* phenotype. The absolute amounts of normal and intron-3-retained *IIL1* transcripts were measured in each single leaf. Single-leaf samples exhibiting different phenotypic statuses were separately collected from three individual plants of the *IIL1-(TTC)_100_/Col-0* lines. The samples included the first normal leaves, the fifth transitional *iil* leaves, and the ninth *iil* leaves (Figure 4A). In parallel, leaves of the same leaf stage were collected from three individual plants of the *IIL1-(TTC)_100_/Col-0* lines that did not show the *iil* phenotype (Figure 4B), as well as from the Col-0 WT that was used as the control (Figure 4C). The absolute amounts of the *IIL1* transcripts in these single-leaf samples were analyzed using quantitative real-time reverse transcription-PCR (qRT-PCR).

**Figure 4.**
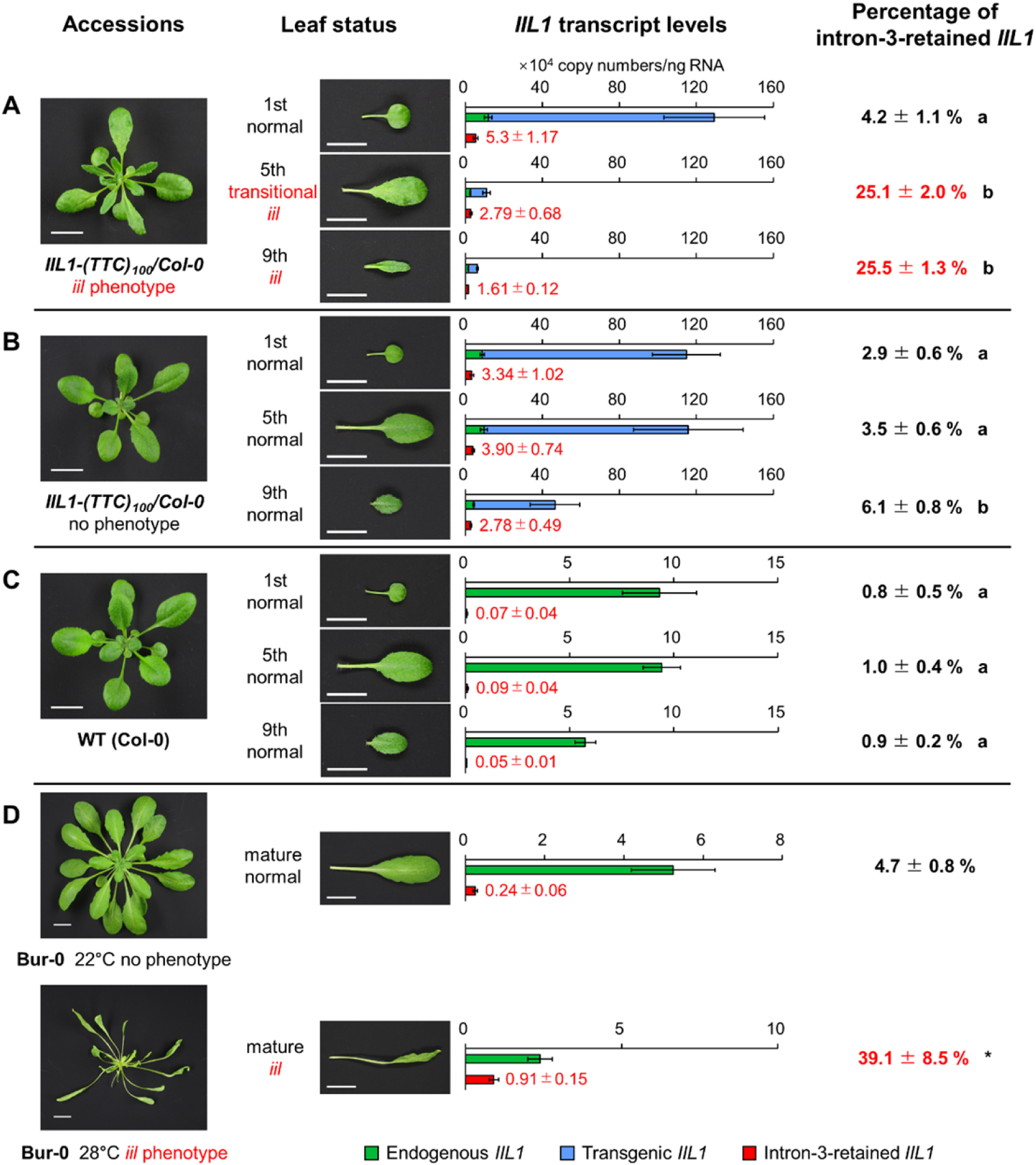
Correlation between the percentage of intron-3-retained *IIL1* transcripts and the occurrence of the *iil* phenotype. (A and B) The first normal leaves, the fifth transitional *iil* leaves, and the ninth *iil* leaves were sampled from four-week-old *IIL1-(TTC)_100_/Col-0* plants showing the *iil* phenotype in the fifth and later leaves (A), and the first, fifth, and ninth normal leaves were sampled from plants that did not show the *iil* phenotype (B). Three single-leaf samples were collected for each kind of leaf from different individuals. The absolute amounts of the *IIL1* transcripts were analyzed in each single-leaf sample by quantitative real-time reverse transcription-PCR (qRT-PCR), using primers 12 and 13 to detect transcripts from transgenic *IIL1* (*IIL1^Bur-0^*allele; blue bars), primers 11 and 13 to detect transcripts from endogenous *IIL1* (*IIL1^Col-0^* allele; green bars), and primers 14 and 15 (Table S1; Figure S10) to detect intron-3-retained *IIL1* transcripts (red bars). The percentage of intron-3-retained *IIL1* transcripts was calculated as “intron-3-retained / (transgenic + endogenous)” *IIL1* transcripts in each sample, and the average and standard deviations are presented. (C) The first, fifth, and ninth normal leaves were sampled from Col-0 wild type. Three single-leaves were collected for each kind of leaf. The absolute amounts of *IIL1* transcripts were analyzed in each single-leaf sample by qRT-PCR, using primers 11 and 13 to detect the transcripts from endogenous *IIL1* (*IIL1^Col-0^* allele; green bars), and primers 14 and 15 to detect intron-3-retained *IIL1* transcripts (red bars). The percentage of intron-3-retained *IIL1* transcripts was calculated as “intron-3-retained / endogenous” *IIL1* transcripts in each sample, and average and standard deviations are presented. (D) The normal and *iil* leaves were sampled from the rosettes of the Bur-0 wild type cultured under 22°C and 28°C, respectively. Three single-leaves were collected for each kind of leaf. The absolute amounts of *IIL1* transcripts were analyzed in each single-leaf sample by qRT-PCR, using primers 12 and 13 to detect transcripts of the endogenous *IIL1* (*IIL1^Bur-0^* allele; green bars), and primers 14 and 15 to detect intron-3-retained *IIL1* transcripts (red bars). The percentage of intron-3-retained *IIL1* transcripts was calculated as “intron-3-retained / endogenous” *IIL1* transcripts in each sample, and the average and standard deviations are presented. One-way analysis of variance (ANOVA) with Duncan’s test was performed within (A, B, and C) groups, and different letters within a group represent the statistical difference (*p* < 0.05). A Student’s *t*-test was performed between mature normal and *iil* leaves of Bur-0 in (D) group. The symbol * indicates statistical difference at *p* < 0.05. Scale bars represent 1 cm.

The genome of the *IIL1-(TTC)_100_/Col-0* lines harbored both transgenic *IIL1* genes derived from Bur-0 (hereafter *IIL^Bur-0^*) and endogenous *IIL1^Col-0^*. We designed allele-specific primers based on the previously described single-nucleotide polymorphisms to separately detect the transcripts from the endogenous (*IIL1^Col-0^* allele) and transgenic (*IIL1^Bur-0^* allele) *IIL1* genes (Figure S10A and B). This enabled us to evaluate the amount of Bur-0 type or Col-0 type *IIL1* transcripts.

In the first normal leaves of the *IIL1-(TTC)_100_/Col-0* lines, the transgenic *IIL1* gene expression was approximately 10-times higher than that of the endogenous *IIL1* gene (Figure 4A; Figure S11A). When the *IIL1-(TTC)_100_/Col-0* lines showed the *iil* phenotype, the expression levels of the transgenic and endogenous *IIL1* were reduced to 8% and 21% on average in the fifth transitional *iil* leaves, respectively; and further decreased to 4% and 13% in the ninth *iil* leaves, respectively (Figure 4A; Figure S11A). This suggested the co-suppression of transgenic and endogenous *IIL1*s in the *IIL1-(TTC)_100_/Col-0* lines exhibiting the *iil* phenotype. By contrast, when the *IIL1-(TTC)_100_/Col-0* lines did not show the *iil* phenotype (Figure 4B and Figure S11B), the expression levels of the transgenic and endogenous *IIL1*s were not reduced in the corresponding fifth normal leaves and were only mildly reduced to approximately 40%∼50% in the ninth normal leaves.

The intron-3-retained *IIL1* transcripts were then quantified using the primer pair annealing to the regions in intron-3 and exon-4 (Figure S10C). In *IIL1-(TTC)_100_/Col-0* lines, no matter whether the *iil* phenotype occurred, the expression levels of intron-3-retained *IIL1* transcripts were higher than those in the Col-0 WT (Figure 4A-C). This suggested that the occurrence of the *iil* phenotype was not attributed to the absolute amount of intron-3-retained *IIL1* transcript. Then we detected the percentage of intron-3-retained *IIL1* transcripts, calculated as “intron-3-retained transcripts / transcripts from transgenic and endogenous *IIL1*s” for each leaf sample. In the Col-0 WT, the percentage of intron-3-retained *IIL1* transcripts was constantly less than 1% in all leaf samples (Figure 4C). In *IIL1-(TTC)_100_/Col-0* lines exhibiting the *iil* phenotype, the percentage of intron-3-retained *IIL1* transcripts was maintained at relatively low levels (4.2%) in the first normal leaves, but strongly increased to approximately 25% in both the fifth transitional and ninth *iil* leaves (Figure 4A). By contrast, when the *IIL1-(TTC)_100_/Col-0* lines did not show the *iil* phenotype, all normal leaf samples maintained relatively low levels of the intron-3-retained *IIL1* transcript (< 6%) (Figure 4B). Therefore, we hypothesized that the increased percentage of intron-3-retained *IIL1* transcripts harboring the TTC repeat lead to the occurrence of the *iil* phenotype. Notably, although the total expression of *IIL1* was drastically decreased in the *iil* leaves, it was still quantitatively equivalent to the levels in the WT (Figure S12). Combined with the observation that the down-regulation of *IIL1* in its T-DNA insertional mutants (*iil1-1* and *iil1-2*) did not result in the *iil* phenotype (Figure S2), our results indicated that the *iil* phenotype was caused by the decrease in absolute levels of total *IIL1* transcripts and the consequent increase in the relative amount of intron-3-retained *IIL1* transcripts harboring the TTC repeat.

To confirm that this was also the case for Bur-0, the transcript levels were quantified from its mature normal leaves and mature *iil* leaves cultured at both 22°C and 28°C, respectively. As shown in Figure 4D, the percentage of the intron-3-retained *IIL1* transcripts was 4.7% in normal leaves, and dramatically increased to 39.1% in the *iil* leaves. These results indicated that the observations for the *IIL1-(TTC)_100_/Col-0* lines showed that the molecular phenomenon also occurred in Bur-0 at elevated temperatures.

### Suppression of the *iil* penetrance by increasing the normal *IIL1* transcripts

Although the increased percentage of intron-3-retained *IIL1* transcripts triggered the *iil* phenotype in individual leaves, it was not clear whether the percentage affected the penetrance of the *iil* phenotype in a whole plant level. Therefore, we examined whether the *iil* penetrance would be altered by changing the percentage of intron-3-retained *IIL1* transcripts in *IIL1-(TTC)_100_/Col-0* lines. The coding sequence (CDS) of the *IIL1* gene (*IIL1-CDS*) without any intron or TTC repeat was cloned from the Bur-0 cDNA, introduced into Col-0 under the control of the *CaMV 35S* promoter, and then introduced into two independent *IIL1-(TTC)_100_/Col-0* lines (#2 and #4) by genetic crossing. The resulting double homozygous lines in the F3 generation were named *IIL1-CDS × IIL1-(TTC)_100_/Col-0*. Because the normal *IIL1* transcripts were additionally expressed from *IIL1-CDS*, the total *IIL1* transcript levels in the *IIL1-CDS × IIL1-(TTC)_100_/Col-0* lines were higher than those in the *IIL1-(TTC)_100_/Col-0* lines (Figure 5A), resulting in a decreased percentage of intron-3-retained *IIL1* transcripts in the ninth leaves of the *IIL1-CDS × IIL1-(TTC)_100_/Col-0* lines (Figure 5B). The penetrance of the *iil* phenotype was investigated in *IIL1-CDS × IIL1-(TTC)_100_/Col-0* lines (Figure 5C). Compared to the corresponding parent *IIL1-(TTC)_100_/Col-0* lines, the penetrance of the *iil* phenotype in two *IIL1-CDS × IIL1-(TTC)_100_/Col-0* lines was greatly decreased to 2% and 1% at stage 1.03, and 13% and 27% at stage 1.08, respectively. These results indicate that the decrease in the percentage of intron-3-retained *IIL1* transcripts caused by the increase in the expression of normal *IIL1* transcripts suppressed the penetrance of the *iil* phenotype in *IIL1-(TTC)_100_/Col-0* lines.

**Figure 5.**
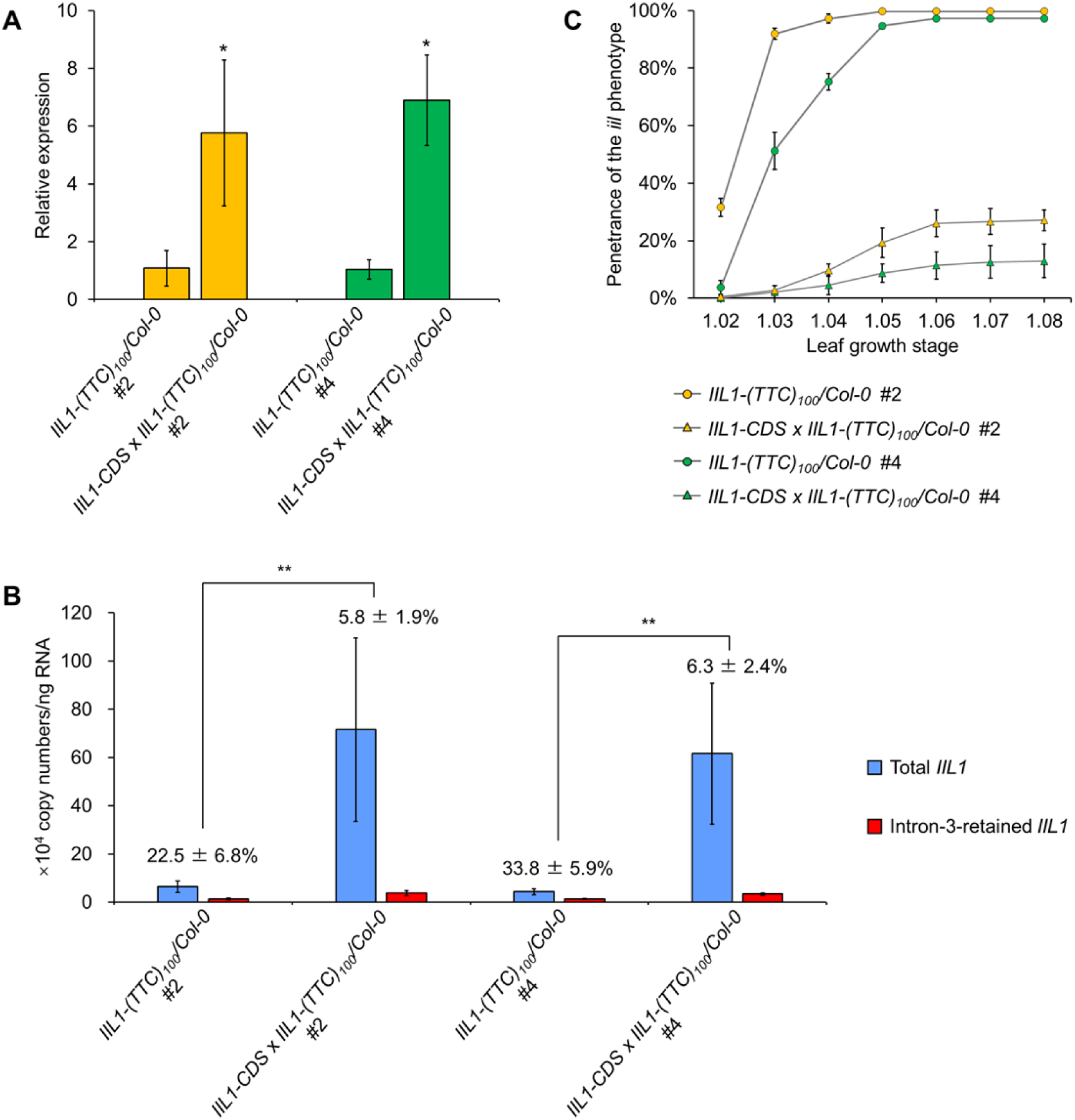
The penetrance of the *iil* phenotype in *IIL1-CDS×IIL1-(TTC)_100_/Col-0* lines. (A) Relative expression of the *IIL1* gene in the *IIL1-CDS×IIL1-(TTC)_100_/Col-0* and corresponding *IIL1-(TTC)_100_/Col-0* parent lines. The relative expression of *IIL1* was measured in eleven-day-old rosettes using primer 9 and 10 (Table S1). Quantitative real-time reverse transcription-PCR was performed with the ΔΔCt method and *UBC9* as the reference gene. The expression levels in the *IIL1-(TTC)_100_/Col-0* lines were set to 1. Data represent means ± standard deviations of three biological replicates. A Student’s *t*-test was performed between the *IIL1-CDS×IIL1-(TTC)_100_/Col-0* and corresponding *IIL1-(TTC)_100_/Col-0* parent lines. The symbol * indicates statistical difference at *p* < 0.05. (B) The percentage of intron-3-retained *IIL1* transcripts in the *IIL1-CDS×IIL1-(TTC)_100_/Col-0* and corresponding *IIL1-(TTC)_100_/Col-0* parent lines. The ninth leaves were sliced from three individuals of four-week-old *IIL1-(TTC)_100_/Col-0* plants showing the *iil* phenotype and phenotypically-normal *IIL1-CDS×IIL1-(TTC)_100_/Col-0* lines. Transcript levels of *IIL1* were analyzed in each single-leaf sample by absolute quantitative real-time reverse transcription-PCR, using primers 9 and 10 (Table S1) to detect the total amount of *IIL1* transcripts, and primers 14 and 15 to detect intron-3-retained *IIL1* transcripts. The percentage of intron-3-retained *IIL1* transcripts was calculated as “intron-3-retained / total” *IIL1* transcripts for each sample. A Student’s *t*-test was performed between the *IIL1-CDS×IIL1-(TTC)_100_/Col-0* and corresponding *IIL1-(TTC)_100_/Col-0* parent lines. The symbol ** indicates statistical difference at *p* < 0.01. (C) The penetrance of the *iil* phenotype in *IIL1-CDS×IIL1-(TTC)_100_/Col-0* and corresponding *IIL1-(TTC)_100_/Col-0* parent lines during leaf development. Data represent means ± standard deviations of three independent experiments. Each experiment was performed using at least 100 individuals for each independent line.

### Suppression of the *iil* phenotype in the *myb28 myb29* background

Finally, we investigated how the intron-3-retained *IIL1* transcripts affected leaf growth. To achieve this, the fact that the IIL1 protein works as a heterodimer was utilized. IIL1 is a large subunit of IPMI that can dimerize with small subunits encoded by three *AtLeuD*s (*AtLeuD1*, *AtLeuD2*, and *AtLeuD3*). The IIL1-AtLeuD3 heterodimer works primarily as an isopropylmalate isomerase and participates in leucine biosynthesis, while the IIL1 heterodimer with either AtLeuD1 or AtLeuD2, functions as a methylthioalkylmalate isomerase in the methionine side-chain elongation pathway and participates in the biosynthesis of AGSLs (Knill *et al*., 2009; Sawada *et al*., 2009; He *et al*., 2010). The expression of *AtLeuD1* and *AtLeuD2*, but not *AtLeuD3*, is positively regulated by the transcription factors MYB28, MYB29, and MYB76 which regulate AGSL biosynthesis (Gigolashvili *et al*., 2007, 2008; Hirai *et al*., 2007; Sønderby *et al*., 2007; Malitsky *et al*., 2008). In our previous study, we have generated a *myb28 myb29* double knockout mutant (*dKO*, hereafter), in which the transcript levels of the *AtLeuD1* and *AtLeuD2* are almost undetectable, while those of *AtLeuD3* are not affected (Li *et al*., 2013). We introduced the *35S_pro_:IIL1-(TTC)_100_* construct into *dKO* to generate *IIL1-(TTC)_100_/dKO* (Figure S3B), and analyzed the penetrance of the *iil* phenotype in five independent lines. Strikingly, the penetrance was extremely low in the *IIL1-(TTC)_100_/dKO* lines (Table 1), indicating the reversion of the *iil* phenotype in this genetic background.

The amounts of the *IIL1* transcripts were then quantified in eight phenotypically-normal individuals randomly selected from independent *IIL1-(TTC)_100_/dKO* lines. Interestingly, some individuals exhibited relatively high percentages for the intron-3-retained *IIL1* transcripts, such as the fifth leaves of three individuals and the ninth leaves of four individuals of the *IIL1-(TTC)_100_/dKO* lines that were higher than 10% (Figure 6B). Even though the percentage of intron-3-retained *IIL1* transcripts in the *dKO* background progressively increased with development and was as high as that in the WT background (Figure 4A), the penetrance of the *iil* phenotype was strongly reduced. This result indicated that the effect of the increased percentage of intron-3-retained *IIL1* transcript was alleviated by the absence of *AtLeuD1* and *AtLeuD2* expression. As the lack of AtLeuD1 and AtLeuD2 proteins presumably promoted the dimerization of IIL1 with AtLeuD3, this further suggested that a certain level of the IIL1-AtLeuD3 heterodimers was required for normal leaf development. In conclusion, increased percentages of intron-3-retained *IIL1* transcripts presumably interfered with the formation of the normal IIL1-AtLeuD3 heterodimers or perturbed the function of normal IIL1-AtLeuD3 heterodimers and resulted in the *iil* phenotype.

**Figure 6.**
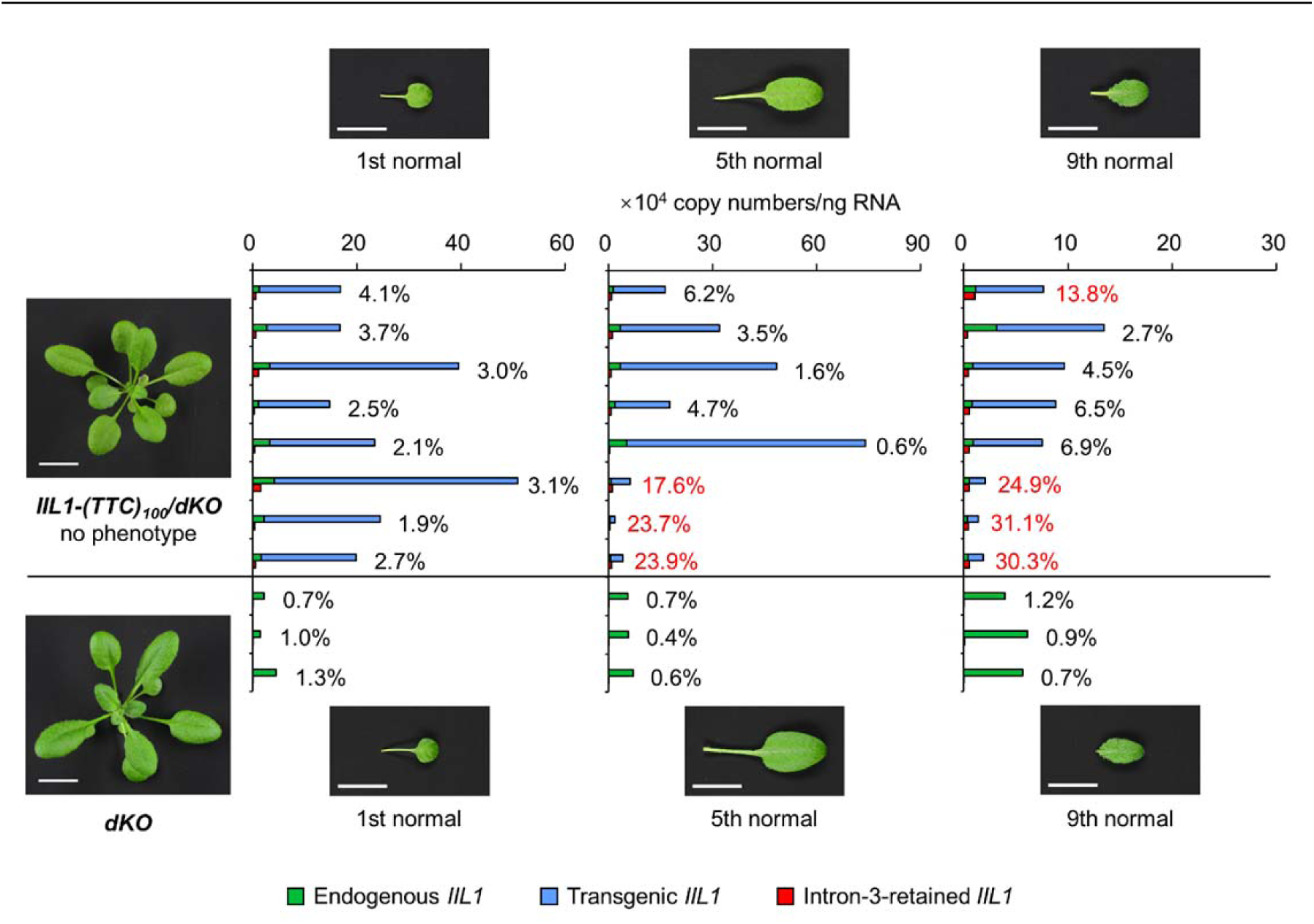
Penetrance of the *iil* phenotype and percentage of intron-3-retained *IIL1* transcripts in the *IIL1-(TTC)_100_/dKO* lines. The first, fifth, and ninth normal leaves were sliced from eight phenotypically-normal individuals randomly selected from different *IIL1-(TTC)_100_/dKO* lines, as well as from three *dKO* individuals. Transcript levels of *IIL1* were analyzed in each single-leaf sample by absolute quantitative real-time reverse transcription-PCR, using primers 12 and 13 (Table S1) to detect the transcripts in the transgenic *IIL1* (*IIL1^Bur-0^* allele), primers 11 and 13 to detect the transcripts in the endogenous *IIL1* (*IIL1^Col-0^*allele), and primers 14 and 15 to detect intron-3-retained *IIL1* transcripts. The percentage of intron-3-retained *IIL1* transcripts was calculated as “intron-3-retained / (transgenic + endogenous)” *IIL1* transcripts for each sample. Values greater than 10% are labelled in red.

## Discussion

### Proposed mechanisms underlying the *iil* phenotype

In this study, we propose an extended scheme for the mechanisms underlying the *iil* phenotype associated with TTC repeats in intron-3 of the *IIL1* gene (Figure 7).

**Figure 7.**
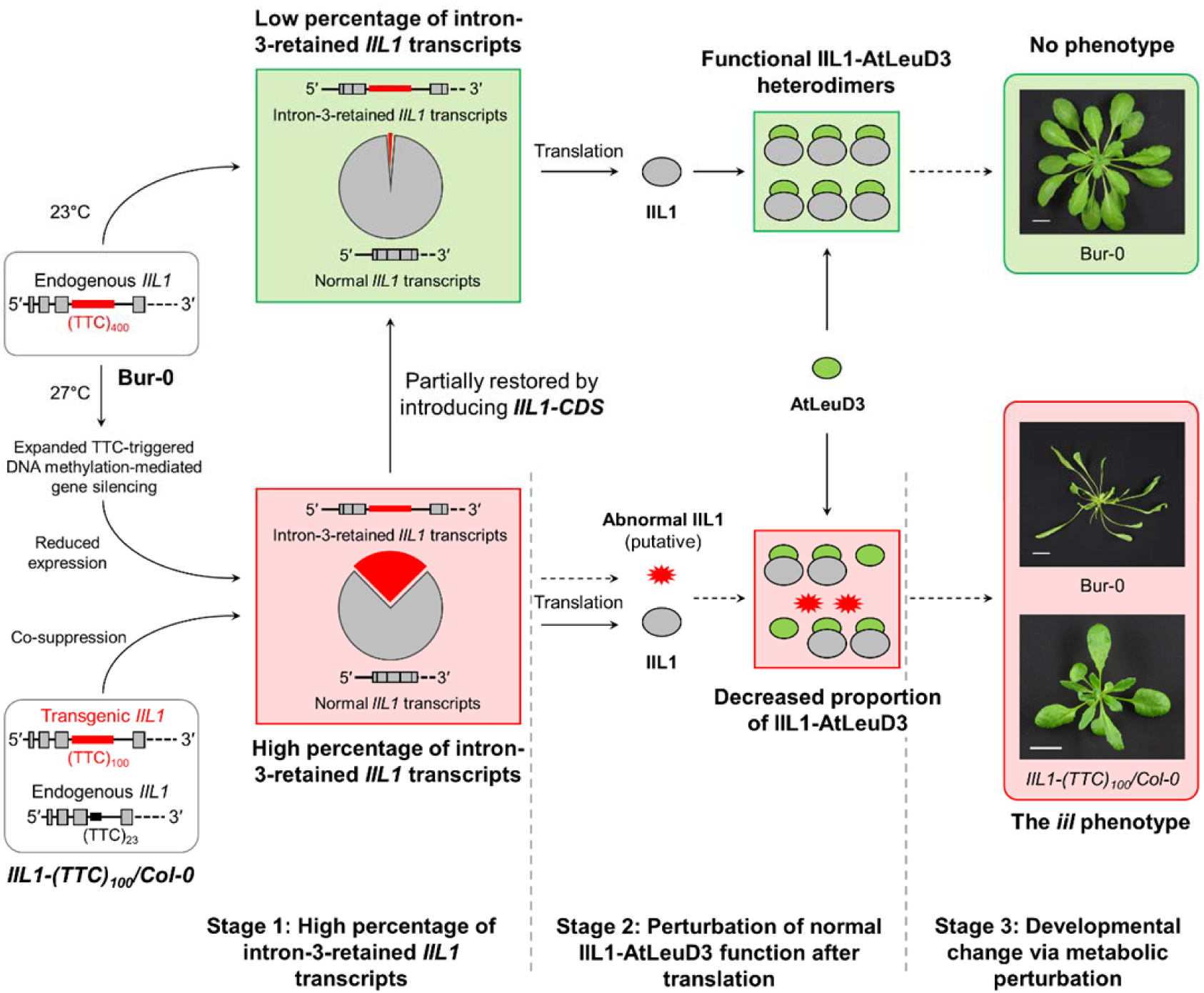
Schematic showing the mechanisms underlying the *iil* phenotype. The percentage of intron-3-retained *IIL1* transcripts is low when Bur-0 is cultured at 23°C, while it is drastically increased when Bur-0 is cultured at 27°C due to expanded TTC-triggered DNA methylation-mediated gene silencing. On the other hand, the percentage of intron-3-retained *IIL1* transcripts is high in the *IIL1-(TTC)_100_/Col-0* lines due to co-suppression. The abnormal IIL1 proteins are putatively translated from intron-3-retained *IIL1* transcripts and interfere with the formation of the normal IIL1-AtLeuD3 heterodimers or perturb the function of the normal IIL1-AtLeuD3 heterodimers, resulting in the *iil* phenotype.

### Stage 1. High percentages of the intron-3-retained IIL1 transcript leads to the iil phenotype

We propose that the occurrence of the *iil* phenotype was caused by increased percentages of the intron-3-retained *IIL1* transcripts that contained the TTC repeat, rather than *IIL1* expression being down-regulated, based on the following observations: (1) The percentage of intron-3-retained *IIL1* transcripts was relatively low in the phenotypically-normal Bur-0, wild-type Col-0, and normal leaves of *IIL1-(TTC)_100_/Col-0*, but was remarkably increased in the Bur-0 showing the *iil* phenotype and *iil* leaves of *IIL1-(TTC)_100_/Col-0*, regardless of the absolute transcript amounts (Figure 4; Figure S12). (2) The penetrance of the *iil* phenotype in *IIL1-(TTC)_100_/Col-0* was repressed by decreasing the percentage of the intron-3-retained *IIL1* transcripts by introducing the *IIL1-CDS* (Figure 5). The previous study also reported that the *iil* phenotype in Bur-0 was suppressed by overexpressing appropriately-spliced mature *IIL1* transcripts (Sureshkumar *et al*., 2009). Once mis-spliced *IIL1* transcripts harboring intron-3 with longer TTC repeats are produced, they might be more tolerant in RNA degradation processes than appropriately-spliced transcripts. A possible explanation for this is that the GAA/UUC repeat enhances RNA stability (Krasilnikova *et al*., 2007). In addition, the content of the retained intron in the transcript regulates the rate of unstable mRNA decay, leading to the enhanced stability of mRNA (Zhao & Hamilton, 2007). A recent study revealed that the degradation of alternatively spliced RNA was differentially affected by the artificial microRNA (amiRNA) in Arabidopsis (Fuchs *et al*., 2021). Given this, the occurrence of the *iil* phenotype by introducing an *IIL1*-targeting amiRNA in the previous study (Sureshkumar *et al*., 2009) might be attributed to the increased percentage of the intron-3-retained *IIL1* transcripts caused by their lower sensitivity to amiRNA-mediated degradation. The proposed mechanisms can explain the absence of the *iil* phenotype in the *IIL1* knockdown mutants *iil1-1* and *iil1-2*, which carry T-DNA insertion in the promoter and 5′-UTR regions of *IIL1*, respectively (Figure S2).

### Stage 2. Intron-3-retained *IIL1* transcript is likely to perturb normal IIL1 function after translation

The next issue raised was how the increased percentage of intron-3-retained *IIL1* transcripts resulted in the *iil* phenotype. In studies of nucleotide repeat diseases in humans, pathogenic mechanisms are thought to be caused by RNA foci formed by repeats-containing mRNA and/or gain-of-function or loss-of-function proteins translated from such mRNAs. For example, GC-rich repeat expansion RNA, such as CCUG in myotonic dystrophy type 2 (Raheem *et al*., 2010; Sznajder *et al*., 2018) and GGGGCC in amyotrophic lateral sclerosis frontotemporal dementia (Niblock *et al*., 2016), forms complicated secondary structures through base-pairing interactions and gives rise to toxic RNA foci (Wojciechowska & Krzyzosiak, 2011; Jain & Vale, 2017). In the case of intron-3-retained *IIL1* transcripts, however, formation of the RNA foci was unlikely, because the UUC repeat expansions did not form RNA secondary structures that caused them (Baralle *et al*., 2008; Ciesiolka *et al*., 2017). Indeed, our *in silico* analysis using the webtools (http://rna.tbi.univie.ac.at/cgi-bin/RNAWebSuite/RNAfold.cgi and https://rna.urmc.rochester.edu/RNAstructureWeb/Servers/Predict1/Predict1.html) predicted that no secondary structure was formed in the region of the TTC repeat in the intron-3 of the *IIL1* transcripts. Otherwise, when intron-retained transcripts are abundantly accumulated and exceed the ability of degradation by the nonsense-mediated RNA decay (Hug *et al*., 2016), they produce truncated proteins. Therefore, the intron-3-retained *IIL1* transcripts, which were abundantly accumulated in the *iil* leaves, might be translated. Sequence analysis showed that intron-3-retained *IIL1* transcripts were predicted to be translated into a truncated N-terminal IIL1 protein with a poly-leucine stretch. Moreover, considering repeat-associated non-AUG translation (*i.e.*, protein translation from expanded repeat-containing transcripts in multiple reading frame without AUG initiation codon) (Pearson, 2011; Zu *et al*., 2011), the intron-3-retained *IIL1* transcripts were predicted to produce truncated N-or C-terminal IIL1 protein with poly-leucine, poly-phenylalanine, or poly-serine stretches. Normal IIL1 protein (*i.e.*, large subunit of IPMI) forms a heterodimer with either of AtLeuD1, AtLeuD2, or AtLeuD3 (*i.e.*, small subunit of IPMI) to function as enzymes. Notably, IIL1-AtLeuD3 is essential for the normal growth and development of plants, since knock-downs of *AtLeuD3* caused growth defect of Arabidopsis (Imhof *et al*., 2014). We proposed that when the percentage of intron-3-retained *IIL1* transcripts was highly increased, the proportion of the abnormal IIL1 protein translated from them to the total IIL1 proteins was increased enough to interfere with the formation of the normal IIL1-AtLeuD3 heterodimer or perturb the function of the normal IIL1-AtLeuD3 heterodimer (Figure 7). This hypothesis was supported by the suppression of the *iil* phenotype: when the appropriately-spliced *IIL1* transcripts were increased by introducing the intron-less *IIL1* CDS, the proportion of functional IIL1-AtLeuD3 was recovered to a sufficient level to support the normal growth and development of the leaves (Figure 5); likewise, suppression of the *iil* phenotype in the *IIL1-(TTC)_100_/dKO* lines (Figure 6) could be explained by assuming that the functional IIL1-AtLeuD3 was increased due to the lack of AtLeuD1 and AtLeuD2.

### Stage 3. Metabolic perturbation could affect leaf development

Because the IIL1-AtLeuD3 heterodimer is the enzyme in leucine biosynthesis and catalyzes the conversion of isopropylmalate, perturbation of normal IIL1 function might affect the metabolism. Indeed, metabolic perturbation was detected in the *iil* leaves of both *IIL1-(TTC)_100_/Col-0* and Bur-0 lines, especially in the accumulation of nitrogen-containing compounds including polyamines, amino acids and their derivatives (Table S2). Nitrogen-containing compounds plays important roles in the developmental control. The examples are found in the gametophore formation in *Physcomitrium patens* specifically affected by arginine metabolism (Kawade *et al*., 2020), the floret development in *Oryza sativa* affected by arginine and ornithine metabolism (Ma *et al*., 2012; Liu *et al*., 2018), stem elongation in Arabidopsis regulated by thermospermine (Kakehi *et al*., 2008), and leaf patterning in Arabidopsis involving γ-aminobutyric acid shunt metabolites (Toyokura *et al*., 2011). Further investigation of metabolic change in the *iil* leaves would lead to identification of the specific metabolic pathway which affects the leaf development.

### TTC repeat length-dependent and progressive occurrence of plant genetic phenotype

In humans, abnormally expanded TNRs lead to a group of inherited neurodegenerative disorders, which are known as trinucleotide repeat expansion diseases (TREDs) (Pearson *et al*., 2005; Mirkin, 2007; La Spada & Taylor, 2010; Pearson, 2011; Martí & Estivill, 2013). TREDs are progressive and TNR length-dependent for onset. Usually, longer TNRs result in a more severe disease status and earlier onset (Walker, 2007; Moseley *et al*., 2006). In contrast to the extensive studies in humans, much less is known about the physiological effects of TNRs on plants and their mechanisms.

Expanded TTC repeats in intron-3 of the *IIL1* gene resulted in an intriguing morphological phenotype of the *IIL1-(TTC)_100_/Col-0* lines, in which the later leaves exhibited severe growth defects while the earlier leaves remained normal (Figure 2C). We also observed obvious heterogeneous rosettes in Bur-0 cultured at 28°C under long-day light conditions (Figure S13). Such heterogenous morphology of the rosettes revealed that the occurrence of the *iil* phenotype was progressive, which is similar to the age-dependent onset of TREDs. In addition, by using *IIL1-(TTC)_100_/Col-0* and *IIL1-(TTC)_23_/Col-0* lines, we demonstrated that the penetrance of the *iil* phenotype was dependent on the TTC repeat length. In the case of most of TREDs, pathogenic TNR sizes range from 40 to several hundred copies. Pathogenic TNR sizes for Friedreich ataxia disease, which is the only reported disorder caused by the GAA/TTC repeat, is more than 200 copies (López Castel *et al*., 2010). In nature, the Col-0 WT, with the endogenous *IIL1* gene and 23 copies of the TTC repeat, did not exhibit the *iil* phenotype even under extreme conditions such as high temperature and UV irradiation (Sureshkumar *et al*., 2009; Tabib *et al*., 2016; Figure S2), because the amount of the intron-3-retained *IIL1* transcripts was quite small in the Col-0 WT (Figure 4C). However, 23 copies of the TTC repeat were enough to cause the *iil* phenotype with relatively low penetrance when it was expressed to an extremely high level when driven by the *CaMV 35S* promoter in the transgenic (*i.e.*, artificial) *IIL1-(TTC)_23_/Col-0* lines (Figure 2E). Therefore, the TTC repeat of the 23 copies seems to be a potential threat to the growth and development of the Col-0 accession.

Based on the similarities between the *iil* phenotype and TREDs, the growth defects caused by the TTC repeat were indeed a kind of “plant genetic disease”. Previous investigations have provided only an example in which TNRs are destructive to plant individuals, given the low biomass and delayed development of individuals exhibiting the *iil* phenotype (Figure 4B). Further research into the mechanisms of the *iil* phenotype is desired not only because it is the first report on the physiological influence of TNR expansions on plants, but also as it provides interesting clues for the pathogenesis of TNR expansion-related diseases.

## Abbreviations

IIL1: ISOPROPYLMALATE ISOMERASE LARGE SUBUNIT1
iil: irregularly impaired leaves
TNR: trinucleotide repeat

## Acknowledgements

We thank Dr. Yuji Sawada for useful discussions.

## Funding

This work was supported by the JSPS Grant-in-Aid for Scientific Research on Innovative Areas, Number JP25113010, to MYH; Number JP25113002 and 19H05672 to HT.

## Author contribution

M.Y.H. and H.T. conceived the project; M.Y.H. supervised experiments; Y.L. and M.Y.H. designed experiments; Y.L., R.L., M.S., A.K., R.S., and A.O. performed experiments; Y.L., K.K., R.S., and A.O. analyzed the data; Y.L. and M.Y.H. wrote the manuscript; Y.L., M.Y.H., H.T., and K.K. revised the manuscript.

## Data availability

Plant materials and source data supporting the findings of this study are available from the corresponding author upon request.

## References

Amorim Franco TM, Blanchard JS. 2017. Bacterial branched-chain amino acid biosynthesis: structures, mechanisms, and drugability. Biochemistry 56: 5849–5865.

Appelhagen I, Lu G, Huep G, Schmelzer E, Weisshaar B, Sagasser M. 2011. TRANSPARENT TESTA1 interacts with R2R3-MYB factors and affects early and late steps of flavonoid biosynthesis in the endothelium of *Arabidopsis thaliana* seeds. The Plant Journal 67: 406–419.

Baralle M, Pastor T, Bussani E, Pagani F. 2008. Influence of Friedreich ataxia GAA noncoding repeat expansions on pre-mRNA processing. American Journal of Human Genetics 83: 77–88.

Binder S. 2010. Branched-Chain Amino Acid Metabolism in *Arabidopsis thaliana*. Arabidopsis Book 8: e0137.

Boyes DC, Zayed AM, Ascenzi R, McCaskill AJ, Hoffman NE, Davis KR, Görlach J. 2001. Growth stage-based phenotypic analysis of Arabidopsis: a model for high throughput functional genomics in plants. The Plant Cell 13: 1499–1510.

Ciesiolka A, Jazurek M, Drazkowska K, Krzyzosiak WJ. 2017. Structural Characteristics of Simple RNA Repeats Associated with Disease and their Deleterious Protein Interactions. Frontiers in Cellular Neuroscience 11: 97.

Eimer H, Sureshkumar S, Singh Yadav A, Kraupner-Taylor C, Bandaranayake C, Seleznev A, Thomason T, Fletcher SJ, Gordon SF, Carroll BJ et al. 2018. RNA-dependent epigenetic silencing directs transcriptional downregulation caused by intronic repeat expansions. Cell 174: 1095–1105.e11.

Fuchs A, Riegler S, Ayatollahi Z, Cavallari N, Giono LE, Nimeth BA, Mutanwad KV, Schweighofer A, Lucyshyn D, Barta A et al. 2021. Targeting alternative splicing by RNAi: from the differential impact on splice variants to triggering artificial pre-mRNA splicing. Nucleic Acids Research 49: 1133–1151.

Gigolashvili T, Yatusevich R, Berger B, Müller C, Flügge UI. 2007. The R2R3-MYB transcription factor HAG1/MYB28 is a regulator of methionine-derived glucosinolate biosynthesis in *Arabidopsis thaliana*. The Plant Journal 51: 247–261.

Gigolashvili T, Engqvist M, Yatusevich R, Müller C, Flügge UI. 2008. HAG2/MYB76 and HAG3/MYB29 exert a specific and coordinated control on the regulation of aliphatic glucosinolate biosynthesis in *Arabidopsis thaliana*. New Phytologist 177: 627–642.

Gur-Arie R, Cohen CJ, Eitan Y, Shelef L, Hallerman EM, Kashi Y. 2000. Simple sequence repeats in *Escherichia coli*: abundance, distribution, composition, and polymorphism. Genome Research 10: 62–71.

Gruer MJ, Artymiuk PJ, Guest JR. 1997. The aconitase family: three structural variations on a common theme. Trends in Biochemical Sciences 22: 3–6.

He Y, Chen B, Pang Q, Strul JM, Chen S. 2010. Functional specification of Arabidopsis isopropylmalate isomerases in glucosinolate and leucine biosynthesis. Plant and Cell Physiology 51: 1480–1487.

Hirai MY, Sugiyama K, Sawada Y, Tohge T, Obayashi T, Suzuki A, Araki R, Sakurai N, Suzuki H, Aoki K et al. 2007. Omics-based identification of Arabidopsis Myb transcription factors regulating aliphatic glucosinolate biosynthesis. *Proceedings of the National Academy of Sciences*, USA 104: 6478–6483.

Hug N, Longman D, Cáceres JF. 2016. Mechanism and regulation of the nonsense-mediated decay pathway. Nucleic Acids Research 44: 1483–1495.

Imhof J, Huber F, Reichelt M, Gershenzon J, Wiegreffe C, Lächler K, Binder S. 2014. The small subunit 1 of the Arabidopsis isopropylmalate isomerase is required for normal growth and development and the early stages of glucosinolate formation. PLoS ONE 9: e91071.

Jain A, Vale RD. 2017. RNA phase transitions in repeat expansion disorders. Nature 546: 243–247.

Kakehi J, Kuwashiro Y, Niitsu M, Takahashi T. 2008. Thermospermine is required for stem elongation in *Arabidopsis thaliana*. Plant and Cell Physiology 49: 1342–1349.

Kawade K, Horiguchi G, Hirose Y, Oikawa A, Hirai MY, Saito K, Fujita T, Tsukaya H. 2020. Metabolic control of gametophore shoot formation through arginine in the moss *Physcomitrium patens*. Cell Reports 32: 108127.

Knill T, Reichelt M, Paetz C, Gershenzon J, Binder S. 2009. *Arabidopsis thaliana* encodes a bacterial-type heterodimeric isopropylmalate isomerase involved in both Leu biosynthesis and the Met chain elongation pathway of glucosinolate formation. Plant Molecular Biology 71: 227–239.

Krasilnikova MM, Kireeva ML, Petrovic V, Knijnikova N, Kashlev M, Mirkin SM. 2007. Effects of Friedreich’s ataxia (GAA)n*(TTC)n repeats on RNA synthesis and stability. Nucleic Acids Research 35:1075–1084.

La Spada AR, Taylor JP. 2010. Repeat expansion disease: progress and puzzles in disease pathogenesis. Nature Reviews Genetics 11: 247–258.

Li Y, Sawada Y, Hirai A, Sato M, Kuwahara A, Yan X, Hirai MY. 2013. Novel insights into the function of Arabidopsis R2R3-MYB transcription factors regulating aliphatic glucosinolate biosynthesis. Plant and Cell Physiology 54: 1335–1344.

Liu C, Xue Z, Tang D, Shen Y, Shi W, Ren L, Du G, Li Y, Cheng Z. 2018. Ornithine δ-aminotransferase is critical for floret development and seed setting through mediating nitrogen reutilization in rice. The Plant Journal 96: 842–854.

López Castel A, Cleary JD, Pearson CE. 2010. Repeat instability as the basis for human diseases and as a potential target for therapy. Nature Reviews Molecular Cell Biology 11: 165–170.

Ma X, Cheng Z, Qin R, Qiu Y, Heng Y, Yang H, Ren Y, Wang X, Bi J, Ma X et al. 2013. *OsARG* encodes an arginase that plays critical roles in panicle development and grain production in rice. The Plant Journal 73: 190–200.

Malitsky S, Blum E, Less H, Venger I, Elbaz M, Morin S, Eshed Y, Aharoni A. 2008. The transcript and metabolite networks affected by the two clades of Arabidopsis glucosinolate biosynthesis regulators. Plant Physiology 148: 2021–2049.

Martí E, Estivill X. 2013. Small non-coding RNAs add complexity to the RNA pathogenic mechanisms in trinucleotide repeat expansion diseases. Frontiers in Molecular Neuroscience 6: 45.

Mirkin SM. 2007. Expandable DNA repeats and human disease. Nature 447: 932–940.

Morgante M, Hanafey M, Powell W. 2002. Microsatellites are preferentially associated with nonrepetitive DNA in plant genomes. Nature Genetics 30: 194–200.

Moseley ML, Zu T, Ikeda Y, Gao W, Mosemiller AK, Daughters RS, Chen G, Weatherspoon MR, Clark HB, Ebner TJ, et al. 2006. Bidirectional expression of CUG and CAG expansion transcripts and intranuclear polyglutamine inclusions in spinocerebellar ataxia type 8. Nature Genetics 38: 758–769.

Murashige T, Skoog F. 1962. A revised medium for rapid growth and bioassay with tobacco tissue cultures. Physiologia Plantarum 15: 472–497.

Nakagawa T, Kurose T, Hino T, Tanaka K, Kawamukai M, Niwa Y, Toyooka K, Matsuoka K, Jinbo T, Kimura T. 2007. Development of series of gateway binary vectors, pGWBs, for realizing efficient construction of fusion genes for plant transformation. Journal of Bioscience and Bioengineering 104: 34–41.

Niblock M, Smith BN, Lee YB, Sardone V, Topp S, Troakes C, Al-Sarraj S, Leblond CS, Dion PA, Rouleau GA et al. 2016. Retention of hexanucleotide repeat-containing intron in *C9orf72* mRNA: implications for the pathogenesis of ALS/FTD. Acta Neuropathologica Communications 4: 18.

Pearson CE, Nichol Edamura K, Cleary JD. 2005. Repeat instability: mechanisms of dynamic mutations. Nature Reviews Genetics 6: 729–742.

Pearson CE. 2011. Repeat associated non-ATG translation initiation: one DNA, two transcripts, seven reading frames, potentially nine toxic entities! PLoS Genetics. 7: e1002018.

Pollard LM, Chutake YK, Rindler PM, Bidichandani SI. 2007. Deficiency of RecA-dependent RecFOR and RecBCD pathways causes increased instability of the (GAA*TTC)n sequence when GAA is the lagging strand template. Nucleic Acids Research 35: 6884–6894.

Raheem O, Olufemi SE, Bachinski LL, Vihola A, Sirito M, Holmlund-Hampf J, Haapasalo H, Li YP, Udd B, Krahe R. 2010. Mutant (CCTG)n expansion causes abnormal expression of zinc finger protein 9 (ZNF9) in myotonic dystrophy type 2. American Journal of Pathology 177: 3025–3036.

Sanchez-Bermejo E, Zhu W, Tasset C, Eimer H, Sureshkumar S, Singh R, Sundaramoorthi V, Colling L, Balasubramanian S. 2015. Genetic architecture of natural variation in thermal responses of Arabidopsis. Plant Physiology 169: 647–659.

Sawada Y, Kuwahara A, Nagano M, Narisawa T, Sakata A, Saito K, Hirai MY. 2009. Omics-based approaches to methionine side chain elongation in Arabidopsis: characterization of the genes encoding methylthioalkylmalate isomerase and methylthioalkylmalate dehydrogenase. Plant and Cell Physiology 50: 1181–1190.

Sønderby IE, Hansen BG, Bjarnholt N, Ticconi C, Halkier BA, Kliebenstein DJ. 2007. A systems biology approach identifies a R2R3 MYB gene subfamily with distinct and overlapping functions in regulation of aliphatic glucosinolates. PLoS ONE 2: e1322.

Sønderby IE, Burow M, Rowe HC, Kliebenstein DJ, Halkier BA. 2010. A complex interplay of three R2R3 MYB transcription factors determines the profile of aliphatic glucosinolates in Arabidopsis. Plant Physiology 153: 348–363.

Sureshkumar S, Todesco M, Schneeberger K, Harilal R, Balasubramanian S, Weigel D. 2009. A genetic defect caused by a triplet repeat expansion in *Arabidopsis thaliana*. Science 323: 1060–1063.

Sznajder ŁJ, Thomas JD, Carrell EM, Reid T, McFarland KN, Cleary JD, Oliveira R, Nutter CA, Bhatt K, Sobczak K et al. 2018. Intron retention induced by microsatellite expansions as a disease biomarker. *Proceedings of the National Academy of Sciences*, USA 115: 4234–4239.

Tautz D, Trick M, Dover GA. 1986. Cryptic simplicity in DNA is a major source of genetic variation. Nature 322: 652–656.

Tabib A, Vishwanathan S, Seleznev A, McKeown PC, Downing T, Dent C, Sanchez-Bermejo E, Colling L, Spillane C, Balasubramanian S. 2016. A polynucleotide repeat expansion causing temperature-sensitivity persists in wild Irish accessions of *Arabidopsis thaliana*. Frontiers in Plant Science 31: 1311.

Toyokura K, Watanabe K, Oiwaka A, Kusano M, Tameshige T, Tatematsu K, Matsumoto N, Tsugeki R, Saito K, Okada K. 2011. Succinic semialdehyde dehydrogenase is involved in the robust patterning of Arabidopsis leaves along the adaxial-abaxial axis. Plant and Cell Physiology 52: 1340–1353.

Varshney RK, Graner A, Sorrells ME. 2005. Genic microsatellite markers in plants: features and applications. Trends in Biotechnology 23: 48–55.

Walker FO. 2007. Huntington’s disease. Lancet 369: 218–228.

Wojciechowska M, Krzyzosiak WJ. 2011. Cellular toxicity of expanded RNA repeats: focus on RNA foci. Human Molecular Genetics 20: 3811–3821.

Zhao C, Hamilton T. 2007. Introns regulate the rate of unstable mRNA decay. Journal of Biological Chemistry 282: 20230–20237.

Zu T, Gibbens B, Doty NS, Gomes-Pereira M, Huguet A, Stone MD, Margolis J, Peterson M, Markowski TW, Ingram MA et al. 2011. Non-ATG–initiated translation directed by microsatellite expansions. *Proceedings of the National Academy of Sciences*, USA 108: 260–265.

